# Evidence for *in vitro* extensive proliferation of adult hepatocytes and biliary epithelial cells

**DOI:** 10.1101/2023.01.03.522656

**Authors:** Takeshi Katsuda, Jinyang Li, Allyson J Merrell, Jonathan Sussman, Juntaro Matsuzaki, Takahiro Ochiya, Ben Z Stanger

## Abstract

Over the last several years, a method has emerged which endows adult hepatocytes with *in vitro* proliferative capacity, producing chemically-induced liver progenitors (CLiPs). However, a recent study questioned the origin of these cells, suggesting that resident liver progenitor cells, but not hepatocytes, proliferate. Here, we provide lineage tracing-based evidence that adult hepatocytes acquire proliferative capacity *in vitro*. Unexpectedly, we also found that the CLiP method allows biliary epithelial cells to acquire extensive proliferative capacity. Interestingly, after long-term culture, hepatocyte-derived cells (hepCLiPs) and biliary-derived cells (bilCLiPs) become similar in their gene expression patterns, and they both exhibit differentiation capacity to form hepatocyte-like cells. Finally, we provide evidence that hepCLiPs can repopulate chronically injured mouse livers, reinforcing our earlier argument that CLiPs can be a cell source for liver regenerative medicine. Moreover, this study offers bilCLiPs as a potential cell source for liver regenerative medicine.

## Introduction

While adult mature hepatocytes (MHs) have been used clinically for cell transplantation therapy, demonstrating safety and short-term efficacy^1^, they are not expandable *in vitro*, posing a major challenge for the field of liver regenerative medicine and limiting the widespread use of this therapy. Over the last several years, we and others have established a methodology to obtain proliferative progenitor-like cells from MHs without gene manipulation^2–8^, which we have called “chemically-induced liver progenitors (CLiPs)”^2,3^. Because CLiPs can regenerate liver tissue when they are transplanted within injured mouse livers^2–8^, this methodology has demonstrated potential for clinical translation, and is becoming more widely studied.

Despite the promising potential of this methodology, there is active debate surrounding the origin of rat CLiPs.^9^ Our original study^2^ and others^7,8^ provided evidence for the ability to convert mouse MHs to proliferative progenitors using *in vitro* lineage tracing. However, the evidence for this observation in rats has been lacking. Recently, Fu et al.^9^ reported a study challenging our argument that CLiPs are derived from MHs. Based on the observation that lineage-traced rat MHs did not derive highly proliferative cells, the authors concluded that the proliferative cells originate from resident liver progenitor cells (LPCs).

Additionally, it has remained unclear whether biliary epithelial cells (BECs), the other epithelial cell type in the liver, can give rise to proliferative cells by using the same strategy. BECs are considered a crucial building block in liver tissue engineering as they comprise the bile drainage system^10–13^. Thus, if the CLiP technique allows BECs to expand *in vitro*, such cells would be a useful cell source for advanced liver tissue engineering.

In this study, using the same rat lineage tracing system as that demonstrated by Fu et al., we first provide evidence that CLiPs are induced from rat MHs. Then, using a mouse lineage tracing system, we provide evidence that the CLiP strategy enables extensive proliferation of BECs as well, offering a new cell source for liver regenerative medicine.

## Results

### CLiPs can be derived from rat adult hepatocytes

As challenged by Fu et al., whether CLiPs can be derived from rat MHs is a growing controversy in the field. Fu et al. identified two morphologically distinct populations that appear under the stimulation of a combination of small molecule inhibitors called YAC (<u>Y</u>-27632, <u>A</u>-83-01 and <u>C</u>HIR99021), which we previously reported as *in vitro* proliferation factors^2^. One population formed colonies composed of small cells and proliferated efficiently, while the second population consisted of large cells and did not proliferate^9^. Using a recently established rat lineage tracing system (Rosa26-LSL-tdTomato rats^14^), Fu et al. found that tdTomato+ MH-derived cells contributed to the large cell population, but not the small cell population. From these data, Fu et al. concluded that CLiPs are not derived from MHs.

Here, we provide evidence against their conclusion using the same rat system. We injected adult Rosa26-LSL-tdTomato rats with AAV8-TBG-iCre (improved Cre), and harvested MHs by the standard percoll gradient method (**Figure 1A**). Consistent with Fu et al.’s observation, the labeling efficiency in this system was very low (0.05-0.1% of hepatocytes were labeled with tdTomato), thereby making the initial plating density of tdTomato+ MHs very sparse (**Figure 1B**). However, this observation led us to hypothesize that tdTomato+ cell clusters appearing in the later time points should be derived from only a single or a few MHs, which would allow us to evaluate the clonal proliferative capacity of MHs. After culturing the cells in the presence of YAC for approximately two weeks, large tdTomato+ cell clusters with small cell morphologies emerged (**Figure 1B**). This observation was readily apparent for multiple microscopic fields in two independent experiments (**Figure S1**). Thus, although we do not have a specific explanation for the conflict between Fu et al.’s observation and ours, we conclude that MHs can give rise to CLiPs under YAC stimulation.

**Figure 1.**
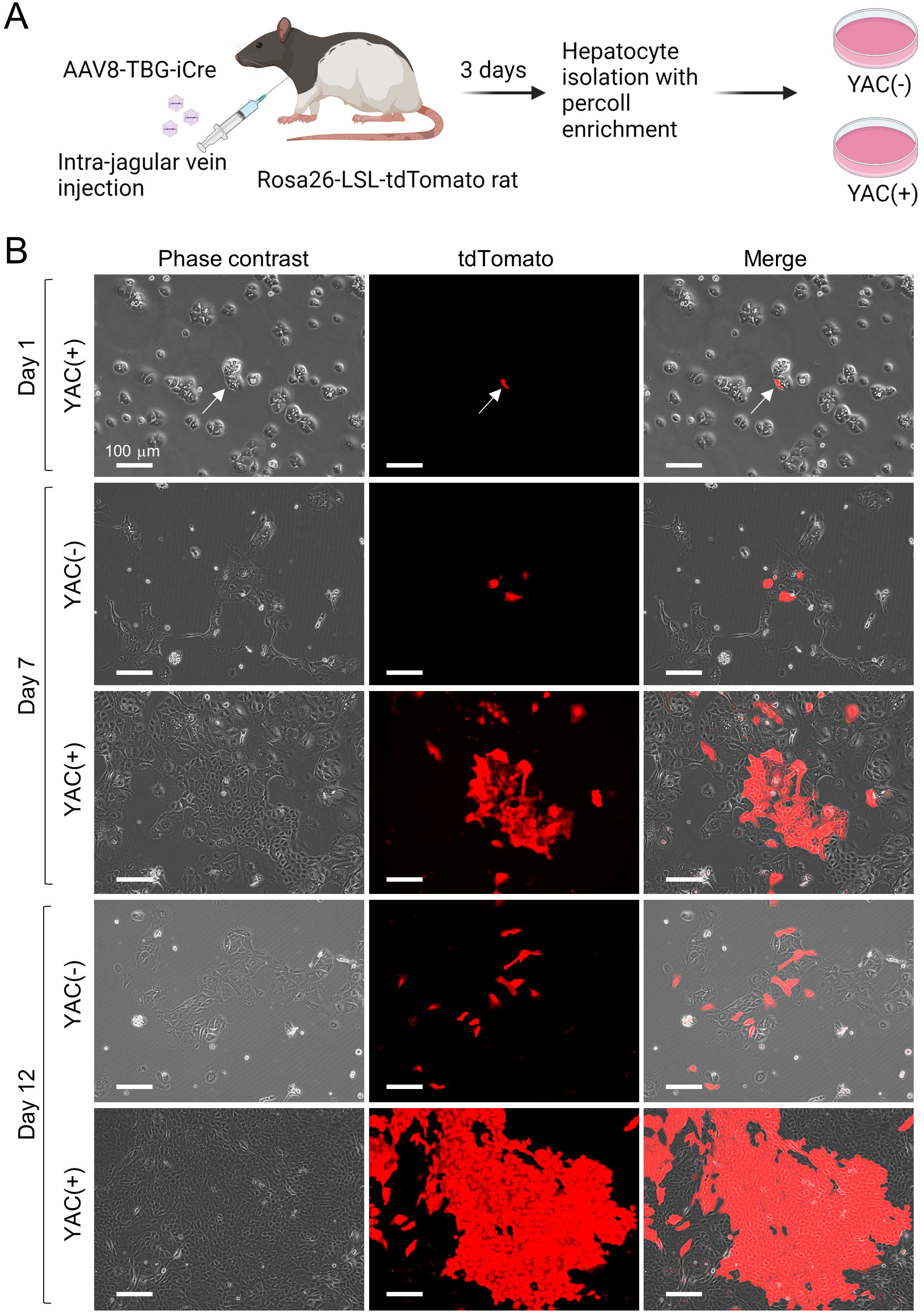
*In vitro* lineage tracing of rat MHs. (A) Schematic representation of *in vitro* lineage tracing of rat MHs. (B) Phase contrast and the corresponding tdTomato fluorescent images of rat hepatocytes cultured with or without YAC at the designated time points. Arrows indicate the single tdTomato-labeled hepatocyte identified at Day 1 in this microscopic field.

### Both hepatocytes and BECs acquire extensive proliferative capacity under YAC culture

Although we and others have provided evidence that hepatocytes can be the origin of CLiPs, it does not rule out the possibility that other cell types also can yield proliferative cells under YAC treatment. Our earlier lineage tracing study in mice was performed during the primary culture period to focus on the emergence of rapidly proliferating cells from hepatocytes^2^. Here, we extended the culture period to test whether hepatocyte-derived cells continue to proliferate after multiple passages and whether non-hepatocyte cells, namely non-parenchymal cells (NPCs), proliferate as well. Using the Rosa26-LSL-EYFP mouse system (**Figure 2A**), we confirmed robust proliferation of YFP+ hepatocyte-derived cells and that these cells exhibited multiple BEC markers, which were not expressed in the original hepatocytes (**Figure 2B**) (note that the term “BEC marker” and “progenitor marker” can be used interchangeably^15^). These BEC markers included those expressed in different stages of hepatobiliary metaplasia *in vivo*^16^ (**Figure S2A**), e.g. Spp1, Sox9 and Hnf1b in the early stage, Itga6 in the intermediate stage, and Krt19 in the late stage. The YFP+ proliferative cells retain expression of Hnf4a, a hepatocyte marker (**Figure 2B**), indicating that they acquired biphenotypic characteristics. We also observed the persistence of *Hnf4a* expression in *in vivo* reprogramming cells (**Figure S2A**), suggesting a partly similar behavior of hepatocyte-derived cells between the *in vivo* reprogramming and the *in vitro* culture.

**Figure 2.**
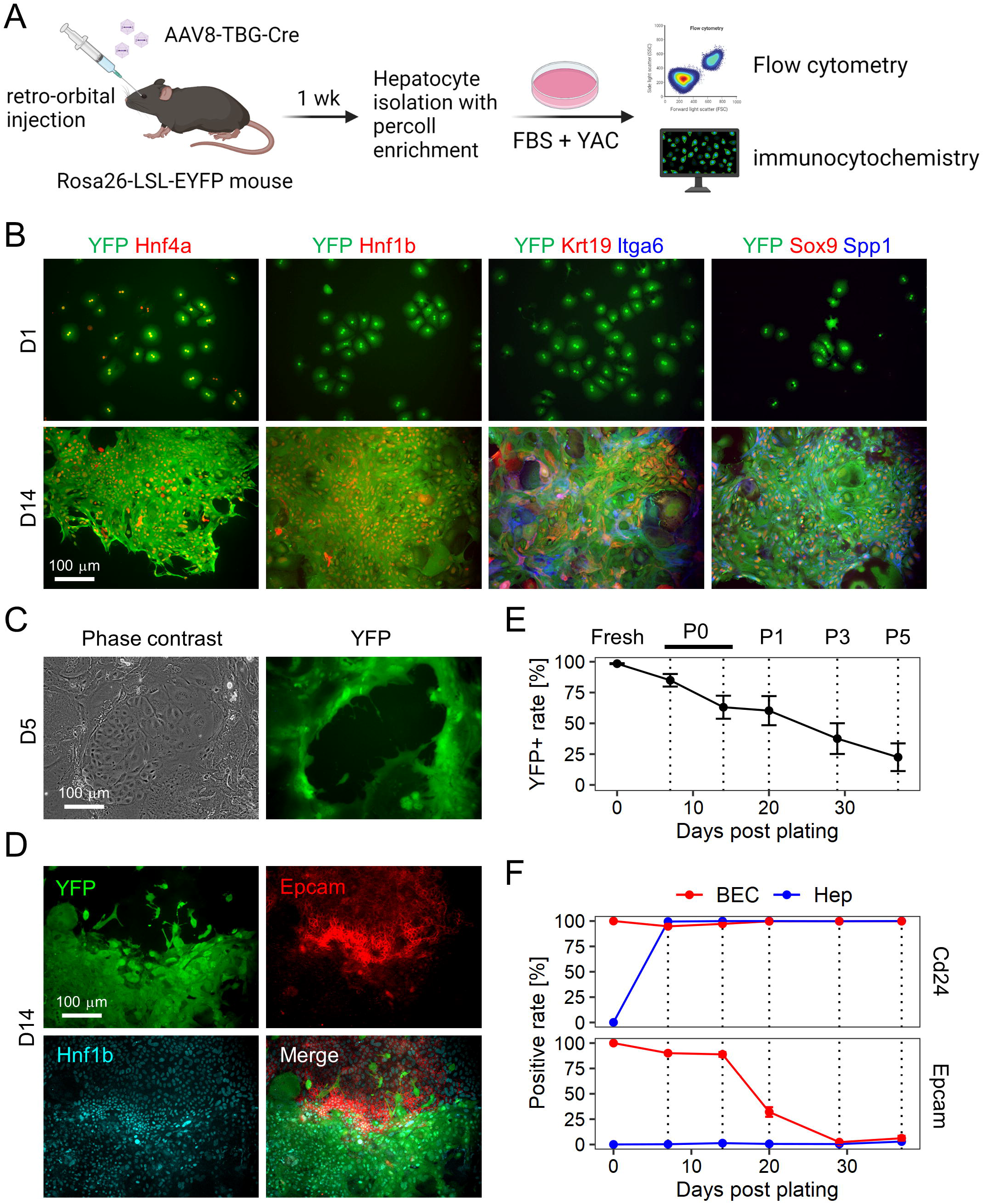
Mouse hepatocytes proliferate and undergo partial biliary reprogramming, while contaminated BECs gradually dominate the population. (A) Schematic representation of *in vitro* lineage tracing of mouse hepatocytes. (B) Confirmation of biphenotypic marker expression of YAC-treated proliferative hepatocytes. (C) Existence of the contaminated BECs becomes evident around 5 days after plating. (D) YFP-epithelial cells express both Hnf1b and Epcam, while YFP+ cells express only Hnf1b. (E) Quantification of YFP+ hepatocyte-derived population by flow cytometry throughout the continued passages. Date represent mean ± SEM (n = 4 donors). (F) Quantification of BEC marker expression in hepatocyte-derived YFP+ cells and that in contaminated YFP-BECs by flow cytometry. Fresh BECs are defined as cells positive for Epcam or Cd24, and the positive values are set to 100%. Date represent mean ± SEM (n = 4 donors).

We then assessed the possibility that NPCs proliferate under the YAC culture condition. Flow cytometry using the percoll gradient-enriched cells yielded a population that was >98% positive for YFP. The ∼2% YFP-negative population contained non-epithelial NPCs, including Cd11b+ Kupffer cells, Cd45+ whole leukocytes, and Cd31+ endothelial cells (**Figure S2B**). Following culture in YAC medium, such non-epithelial NPCs decreased over time and became almost undetectable by the end of primary culture (**Figure S2C**). By contrast, we observed that YFP-epithelial cells grew rapidly in primary culture (**Figure 2C**), and these cells were confirmed to be BECs based on the expression of Hnf1b and Epcam (**Figure 2D**). Considering the rarity of BECs detected in the percoll gradient-enriched population (**Figure S2D**), the growth rate of BECs was much faster than that of hepatocytes. Moreover, BEC proliferation was apparent as early as 2-3 days after plating as assessed by microscopic observation, while hepatocyte proliferation became apparent after around 4-5 days (data not shown). Consequently, the YFP+ cell fraction steadily decreased over the period of the subsequent passages (**Figure 2E**). To assess the biliary characteristics, we monitored the protein expression of two BEC surface markers, Cd24 and Epcam. In a mouse hepatobiliary reprogramming model, we recently found that Cd24 serves as an early-to-intermediate marker of biliary reprogramming of hepatocytes, while Epcam serves as a intermediate-to-late reprogrammed marker (Katsuda et al., *submitted*). Thus, these two markers would give insight into the plastic characteristics of hepatocytes and BECs *in vitro*. We found that Cd24 was readily expressed in all the BECs and hepatocytes after 5 days in primary culture, while Epcam expression was present only in BECs throughout the culture until the fifth passage (P5) (**Figure 2F**). Interestingly, we found that Epcam expression was lost by P3 in BECs, while Cd24 expression was retained in both populations throughout the culture period (**Figure 2F**). These results indicate that hepatocytes partly gained biliary characteristics (Cd24 expression), while BECs partly lost biliary characteristics (Epcam expression). Intriguingly, when sorted BECs were cultured with YAC (**Figure S3A**), they partly exhibited morphologies resembling hepatocytes, as characterized by polygonal and cytoplasm-rich morphologies with occasional binucleation (**Figure S3B**), a characteristic feature of hepatocytes^17^. Collectively, these results suggest that hepatocytes and BECs converge into a similar cell type under YAC culture. Thus, hereafter, we refer to these YAC-induced proliferative cell populations as hepatocyte-derived CLiPs (hepCLiPs) and BEC-derived CliPs (bilCLiPs) respectively. We confirmed that both cell types had almost unlimited proliferative capacity (no reduction in proliferation after at least 20 passages, where cell density was diluted by 1/20 to 1/40 in each passage). These results lead us to modify our earlier argument to conclude that both hepatocytes and BECs can be the origin of CLiPs, not just hepatocytes.

### Hepatocyte-derived and BEC-derived cells become similar *in vitro*

We then sought to characterize hepCLiPs and bilCLiPs in terms of their gene expression profiles. Using a Rosa26-LSL-EYFP mouse, we generated clonal hepCLiP (n = 23) and bilCLiP cell lines (n = 11) (**Figure 3A**). We compared their gene expression patterns along with freshly-isolated hepatocytes and BECs. As shown in **Figure 3B**, hepCLiPs and bilCLiPs demonstrated higher similarity to BECs than to hepatocytes. This is likely due to the decreased expression of most of the assessed hepatocyte genes and the upregulation of BEC genes in hepCLiPs (**Figure 3B**). Some of the assessed BEC genes, including *Tacstd2, Grhl2* and *Epcam*, were substantially decreased in bilCLiPs, but many others, including *Spp1, Hnf1b, Itga6* and *Krt19* were retained throughout the culture period. The expression of these bilCLiP-retained genes was upregulated in hepCLiPs, thereby making their overall expression patterns similar to each other (**Figure 3B**). We also observed that many bilCLiP-retained genes were also acquired at early-to-intermediate stages of hepatobiliary metaplasia *in vivo*; however, many bilCLiP-downregulated genes are acquired at the late stage of hepatobiliary metaplasia with a few notable exceptions such as *Krt19* and *Krt7* (**Figure S2A**).

**Figure 3.**
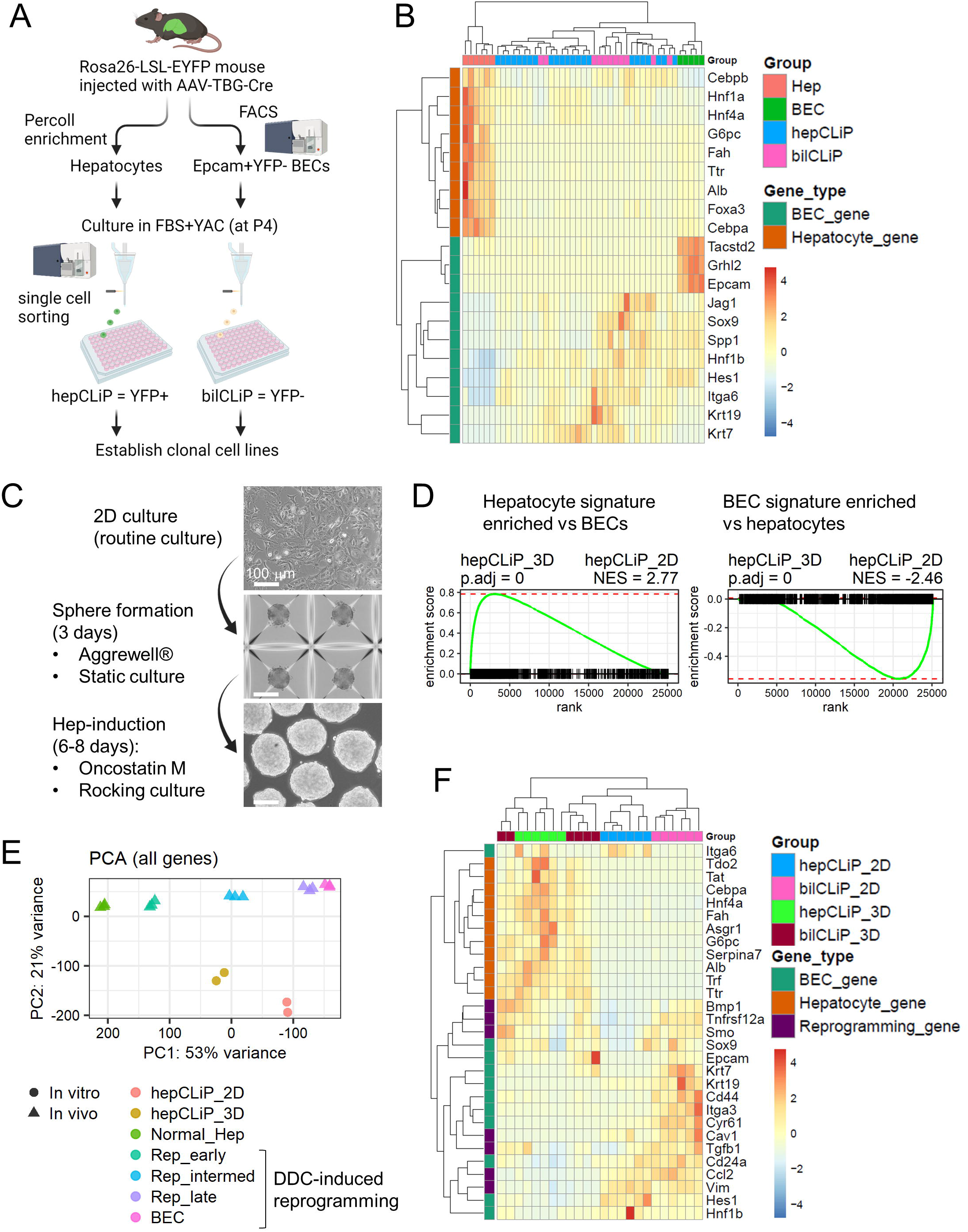
Both hepCLiPs and bilCLiPs exhibit BEC-like phenotypes under 2D culture, while they become more hepatocyte-like under the 3D culture. (A) Schematic of strategy to establish hepCLiPs and bilCLiPs. (B) Heatmap of hepatic and BEC markers as assessed by qRT-PCR with clonal hepCLiPs (n = 23), bilCLiPs (n = 11), fresh hepatocytes (n = 6), and fresh BECs (n = 5). (C) Schematic of 3D culture-based hepatocyte induction of hepCLiPs and bilCLiPs. Images obtained for one of the hepCLiP clones are shown as a representative example. (D) Gene set enrichment analysis (GSEA) comparing hepCLiPs cultured under 2D and 3D conditions using sets of hepatocyte-enriched genes compared with BECs and BEC-enriched genes compared with hepatocytes ^16^ (n = 2 independent donor-derived hepCLiPs). (E) PCA mapping of hepCLiPs cultured in 2D and 3D (n = 2) along with *in vivo* reprogrammed cells (n = 3). *In vivo* samples were harvested from normal mouse livers and those under hepatobiliary reprogramming induced by challenging the mice with 0.1% 3,5-diethoxycarbonyl-1,4-dihydrocollidine (DDC). For *in vivo* reprogramming experiments, the cells were harvested by FACS. (F) Heatmap of hepatic and BEC markers as assessed by qRT-PCR with clonal hepCLiPs (n = 6 clones from 2 donors) and bilCLiPs (n = 6 clones from 1 donor).

While CLiPs resemble BEC-like cells in the routine monolayer culture, they can re-differentiate into hepatocyte-like cells under a condition which induces the hepatocyte phenotype^7,8^. We were interested in whether this is also the case for bilCLiPs. First, using a new culture system (**Figure 3C**) where CLiPs were cultured under a three dimensional (3D) condition in the presence of hepatocyte-inducible cytokine Oncostatin M,^18^ we confirmed that hepCLiPs re-differentiated to hepatocyte-like cells by RNA sequencing (**Figure 3D**). When compared with the *in vivo* reprogramming context, the extent of reprogramming of the 2D-cultured hepCLiPs (hepCLiP_2D) was similar to that of late reprogrammed cells *in vivo*, while that of 3D-cultured hepCLiPs (hepCLiP_3D) reduced to the level of intermediately reprogrammed cells *in vivo* (**Figure 3E**). We then found that bilCLiPs also exhibited differentiation capacity toward hepatocyte-like cells in 3D culture (**Figure 3F**). Under the 2D condition, bilCLiPs (bilCLiP_2D) exhibited relatively more BEC or reprogrammed cell characteristics^19^ than hepCLiP_2D (**Figure 3F**). However, upon culturing in the 3D condition, bilCLiPs (bilCLiP_3D) upregulated the expression of most of the assessed hepatocyte genes. This is further evidenced by the fact that some of the bilCLiP_3D samples were clustered with hepCLiP_3D samples (**Figure 3F**). Thus, these results demonstrate that both hepCLiPs and bilCLiPs behave as BEC-like cells under the 2D culture, while they readily differentiate into hepatocyte-like cells under the 3D culture.

### hepCLiPs regenerate injured mouse livers of FRG mice

Finally, we sought to re-evaluate our earlier argument that hepatocyte-derived CLiPs can regenerate an injured liver, which was not fully proven in our previous study using the rat CLiPs^2^. To this end, we assessed the repopulation capacity of a clonal hepCLiP cell line in FRG (*Fah*^-/-^ *Rag2*^-/-^ *Il2rg*^-/-^) mice, a well-established chronic liver injury model which offers a permissive environment for repopulation by transplanted hepatocytes^20^ (**Figure 4A**). Given that it would be beneficial to know whether there is a difference in the repopulation efficiency between hepCLiP_2D and hepCLiP_3D cells, we prepared cells from both conditions (See Experimental Procedures for details) (**Figure 4A**). At 3.8 months after intrasplenic transplantation, both hepCLiP_2D and hepCLiP_3D cells exhibited robust repopulation in the host livers (**Figure 4B**). The repopulation efficiency was roughly estimated to be comparable between hepCLiP_2D and hepCLiP_3D cells (**Figure 4C**). The sizes of CFP+ repopulated nodules were larger when transplanted with hepCLiP_2D cells, while the number of nodules formed in the FRG livers tended to be great when transplanted by hepCLiP_3D cells (**Figure 4D**). This suggests that hepCLiPs treated with hepatic induction prior to transplantation have higher engraftment efficiency in the host liver compared to hepCLiPs maintained under the routine monolayer culture condition. Since the engraftment efficiency is one of the key determinants for successful cell transplantation therapy^21^, our observation may give an insight into strategies to improve CLiP-based cell transplantation therapy.

**Figure 4.**
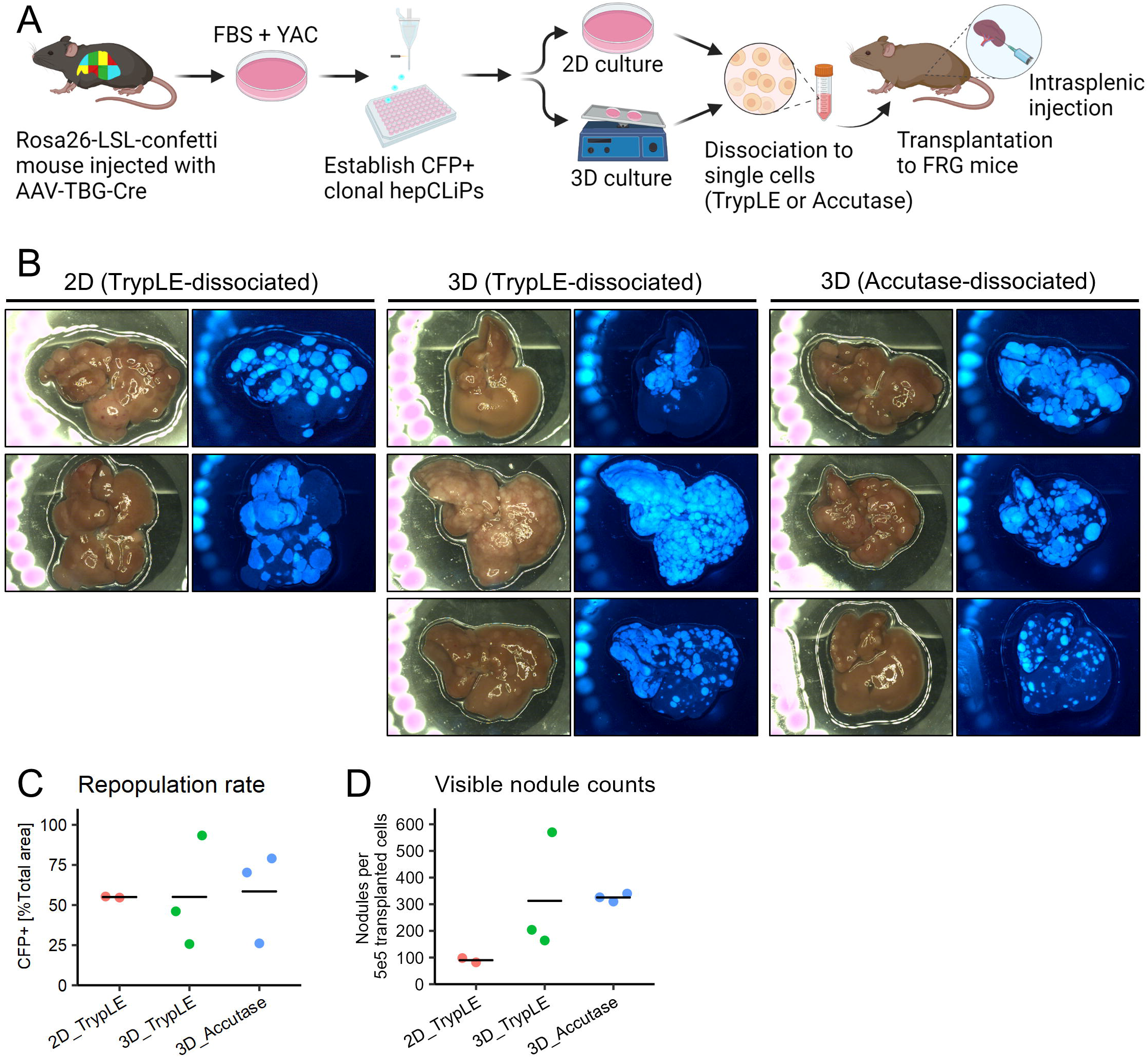
Mouse hepCLiPs repopulate chronically injured mouse livers. (A) Schematic of experimental design for the establishment of CFP-labeled clonal hepCLiPs and the repopulation assay for one of these CFP+ hepCLiP clones using the FRG mouse system. (B) Macroscopic brightfield and fluorescent images of the FRG mouse livers. Livers were harvested from the host mice which were treated with the nitisinone cycle for 3.8 months. (C) Estimation of repopulation efficiency based on the gross fluorescent images described in (B). The horizontal bars indicate the mean values. (D) Number of nodules visible on the liver surface were counted on each image shown in (B) and represented as “per transplanted 5 × 10^5^ cells” (note that 5 × 10^5^ cells/mouse were transplanted for 2D_TrypLE and 3D_TrypLE groups, while 2.5 × 10^5^ cells/mouse were transplanted for 3D_Accutase. See Experimental Procedures for details). The horizontal bars indicate the mean values.

## Discussion

This study provides lineage tracing-based evidence that hepatocytes can be expanded *in vitro* using the small inhibitor cocktail YAC both in rats and mice. Transplantation experiments further confirmed that hepatocyte-derived CLiPs regenerate the mouse liver with chronic injury, thereby providing evidence that hepatocytes can be a cell source for liver regenerative medicine. We also provided evidence that BECs can acquire proliferative capacity in the presence of YAC and exhibit a similar gene expression pattern as that of hepCLiPs. Moreover, bilCLiPs exhibit the capacity to differentiate to hepatocyte-like cells. Future studies are needed to further characterize bilCLiPs, including their contribution to regeneration of the liver parenchymal tissue and bile ducts in the relevant mouse injury models.

In addition to the new data provided above, we delineate possible conflicts between Fu et al.’s interpretation and our own^9^. The term “LPC” is used in Fu et al.’s study to refer to fetal hepatoblasts which persist in the adult liver. However, the authors have not provided evidence for the existence of such cells in adult rats. Although not stated explicitly in their paper, it is possible that LPCs refer to “small hepatocytes,” which are known as a subset of rat MHs with the capacity for several rounds of cell cycle *in vitro*^22^. Due to their small sizes, these cells can be enriched in the supernatant following low-speed centrifugation^22^, as Fu et al. also demonstrated. Small hepatocytes are indistinguishable from MHs except for their size difference and that they start to express Cd44 in culture^23^. Thus, it is more reasonable to regard small hepatocytes as a subset of MHs rather than resident LPCs. After carefully examining Fu et al.’s report, we propose that it is more likely that the “small cell population” they identified is identical to the reprogrammable 2c subset of MHs, which we identified as the origin of CLiPs^2^.

In our follow-up study, we further characterized the difference of 2c, 4c, and 8c populations of rat MHs and identified a small subset of 2c MHs which exhibit progenitor-like features, as characterized by weak expression of *Epcam, Prom1*, and *Lgr5*^24^. This implies that this 2c subpopulation may express progenitor markers more readily in culture than the rest of MHs, which could explain the heterogeneity Fu et al. observed. We emphasize that these cells are not resident LPCs, since the overall gene expression pattern of these cells before culture was very similar to that of 4c and 8c hepatocytes^24^. Moreover, the 2c subpopulation was clearly distinct from BECs^24^, which are known to share more features with LPCs at the transcriptome level^2^. Notably, we reported that the diameters of 2c, 4c and 8c MHs were 18.0 ± 3.3 μm, 24.4 ± 3.6 μm and 29.3 ± 4.3 μm, respectively (mean ± S.D.)^24^, which was reasonably consistent with Fu et al.’s observation that the diameter of purified small cells was 16.8 μm on average, while that of MHs was 24.5 μm^9^. Given that the remainder of Fu et al.’s study described very similar observations as to what we reported previously, namely the demonstration of hepatic/biliary differentiation capacities^2^, we would consider that their report does not challenge our findings, but rather faithfully reproduces our earlier study, thereby reinforcing the concept of the CLiP methodology.

## Acknowledgements

We thank the Penn Xenograft Core for providing the FRG mice, performing transplantation experiments, and taking care of the nitisinone cycles to maintain the mice. We thank Dr. Bin Li (Grompe Lab in OHSU) for providing valuable advice on the repopulation experiments. This work was supported by NIH grants R01DK083355, the Fred and Suzanne Biesecker Pediatric Liver Center, the Abramson Family Cancer Research Institute, the Mochida Memorial Foundation for Medical and Pharmaceutical Research, the Uehara Memorial Foundation, the Kanae Foundation, and the Osamu Hayaishi Memorial Scholarship for study abroad of the Japanese Biochemical Society.

## Competing interests

None declared.

## Author contributions

Conceptualization, T.K.; Experiments, T.K., A.J.M.; Data Analysis, T.K., J.L, J.S.; Investigation, T.K., A.J.M., J.S., J.M.; writing, T.K., J.S.; Funding Acquisition, T.K., B.Z.S., T.O.

## Experimental Procedures

### Rats

Rosa26-LSL-tdTomato rats on Long Evans background were injected with AAV8-TBG-iCre (improved Cre) virus (VectorBio Lab) via the jugular vein. Three days following injection, hepatocytes were harvested as described below. Studies were conducted in accordance with the guidelines of the Institute for Laboratory Animal Research, National Cancer Center Research Institute.

### Mice

Heterozygous or homozygous Rosa26-LSL-EYFP and Rosa26-LSL-Confetti mice on C57BL/6 backgrounds at 6-10 weeks of age were injected retro-orbitally with 2.5 × 10^11^ viral particles of AAV8-TBG-Cre virus (Penn Vector Core). One week following injection, liver cells were isolated as described below. FRG mice^20^ at 5-6 months of age were used as host animals for the repopulation assay. Hepatobiliary reprogramming was modelled in Rosa26-LSL-Cas9-IRES-EGFP mice on C57BL/6 by challenging the mice with 0.1% 3,5-diethoxycarbonyl-1,4-dihydrocollidine (DDC) diet (Envigo) 1 week after AAV8-TBG-Cre injection. Studies were conducted in accordance with the NIH and University of Pennsylvania Institutional Animal Care and Use Committee guidelines.

### Cell isolation from rats

A step-by-step protocol is available online^25^. Briefly, the liver was digested by two-step collagenase perfusion. The extracted liver was further digested in a collagenase solution at 37 °C for 15 min. The digested liver was then filtered twice through a sterilized cotton mesh, and the cells were collected via centrifugation at 57 ×*g* for 1 min. Following resuspension in 50 ml/tube of E-MEM (Sigma), large cell aggregates were eliminated by filtering the cell suspension with a 60-μm stainless cell strainer (Ikemoto Scientific Technology), and the cells were collected by centrifugation at 57 ×*g* for 1 min. Then, the cells were resuspended in 24.5 ml/tube of complete Percoll media (composed of 25 ml of L-15 medium [Life Technologies] supplemented with 0.429 g/L HEPES [Sigma], 2 g/L BSA [Sigma], 1 × 10^−7^ M insulin [Sigma], 2.4 ml of 10× HBSS(-) [Life Technologies], and 21.6 ml of Percoll [GE Healthcare]). Dead cells were removed via centrifugation at 57 ×*g* for 10 min. Then, the cells were washed in 50 ml/tube of E-MEM twice via centrifugation at 57 ×*g* for 2 min, and used for cell culture.

### Cell isolation from mice

Livers were perfused with 40 ml of 1× HBSS, followed by 40 ml HBSS/1 mM EGTA, and 40 ml of HBSS/5 mM CaCl_2_/40 µg/ml liberase. Following perfusion, livers were mechanically dissociated with tweezers, resuspended in 10 ml DMEM supplemented with 5% fetal bovine serum (FBS) (hereafter called DMEM5), and filtrated with a 70 μm cell strainer. The cells were centrifuged at 50 ×*g* for 5 minutes. Then, the cells were resuspended in complete percoll solution (10.8 ml percoll, 12.5 ml DMEM5 and 1.2 ml 10× HBSS(-)), and centrifuged at 50 ×*g* for 10 minutes. The cells were washed in 10 ml DMEM5 once by centrifugation at 50 ×*g* for 5 minutes, then the cells were used for cell culture, flow cytometry, FACS and RNA extraction.

### Cell culture

For rat cell culture, FBS-free small hepatocyte culture medium (SHM) was used as basal medium^26^, namely, DMEM/F12 (Corning) containing 2.4 g/L NaHCO3 and L-glutamine, which was supplemented with 5 mM HEPES (Corning), 30 mg/L L-proline (Sigma), 0.05% BSA (Sigma), 10 ng/ml epidermal growth factor (Sigma), insulin-transferrin-serine (ITS)-X (Thermo), 10^−7^ M dexamethasone (Dex) (Sigma), 10 mM nicotinamide (Sigma), 1 mM ascorbic acid-2 phosphate (Wako), and 1× antibiotics (anti-anti (Thermo) or gentamycin (Gemini Bio-Products)). Cells were cultured in SHM with or without 10 mM Y-27632 (Wako), 0.5 mM A-83-01 (Wako), and 3 mM CHIR99021 (Axon Medchem). For mouse cell culture, we used SHM supplemented with 10% FBS and YAC. Isolated cells were suspended in SHM, SHM+YAC, or SHM+10%FBS+YAC at 1 × 10^4^ cells/ml, and plated on collagen-coated plates (IWAKI) according to the surface areas (0.5 ml/well, 1 ml/well, 2 ml/well and 10 ml/plate for a 24 well-, 12 well-, 6 well- and 10 cm plate respectively). Culture medium was replaced every 2-3 days. When cells reached 70-100% confluence, they were harvested by standard trypsinization, and passaged to new plates at 1/20-1/40 dilution.

## Notes

### Competing Interest Statement

The authors have declared no competing interest.

