## Supplemental information for "Evidence for *in vitro* extensive proliferation of adult hepatocytes and biliary epithelial cells"

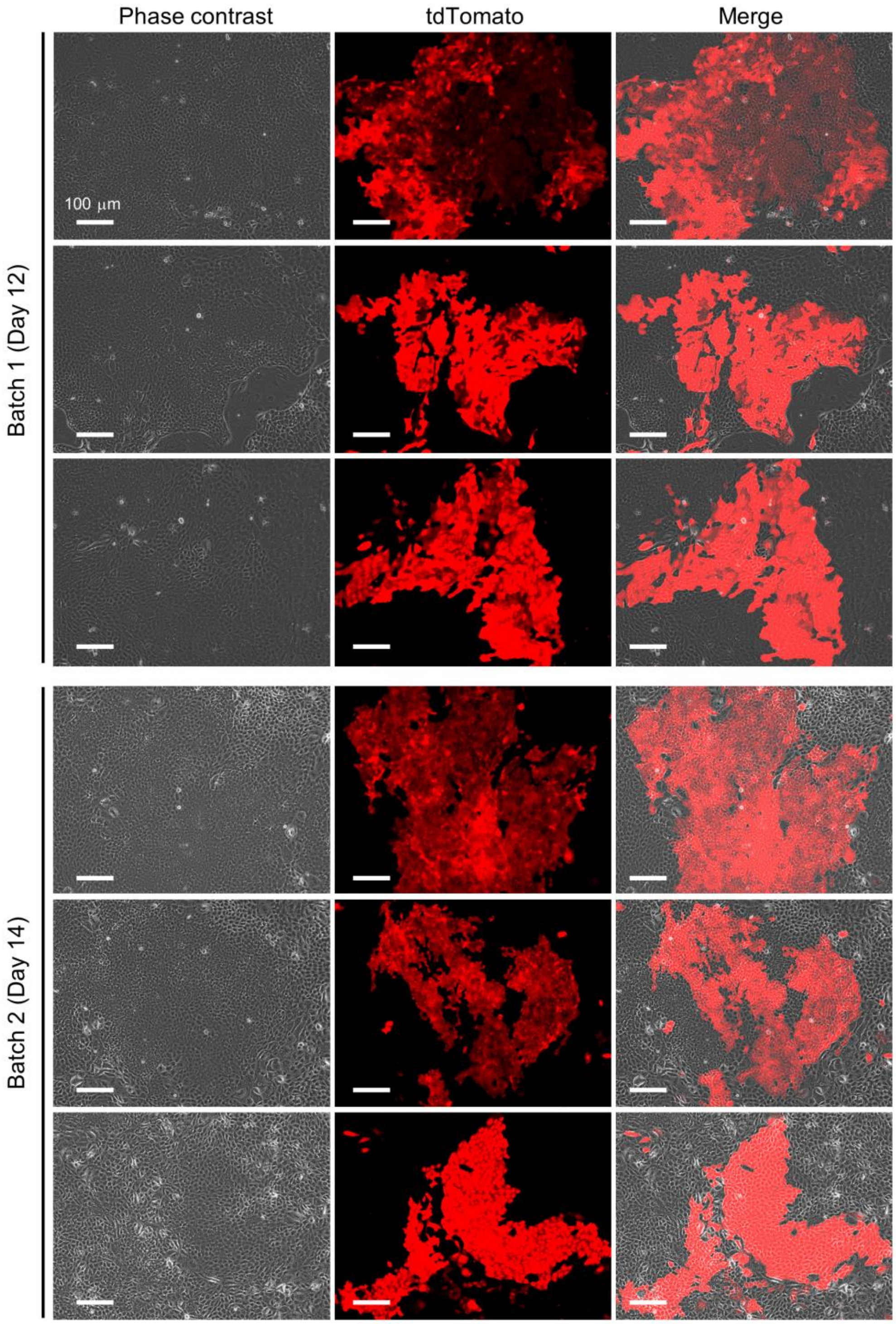


**Figure S1. Evidence for robust expansion of rat hepatocyte-derived cells.**
Additional data to support the observation in **Figure 1B**. The data were obtained from two independent experiments. Three representative fields are shown for each batch where tdTomato+ cells formed a large cluster.

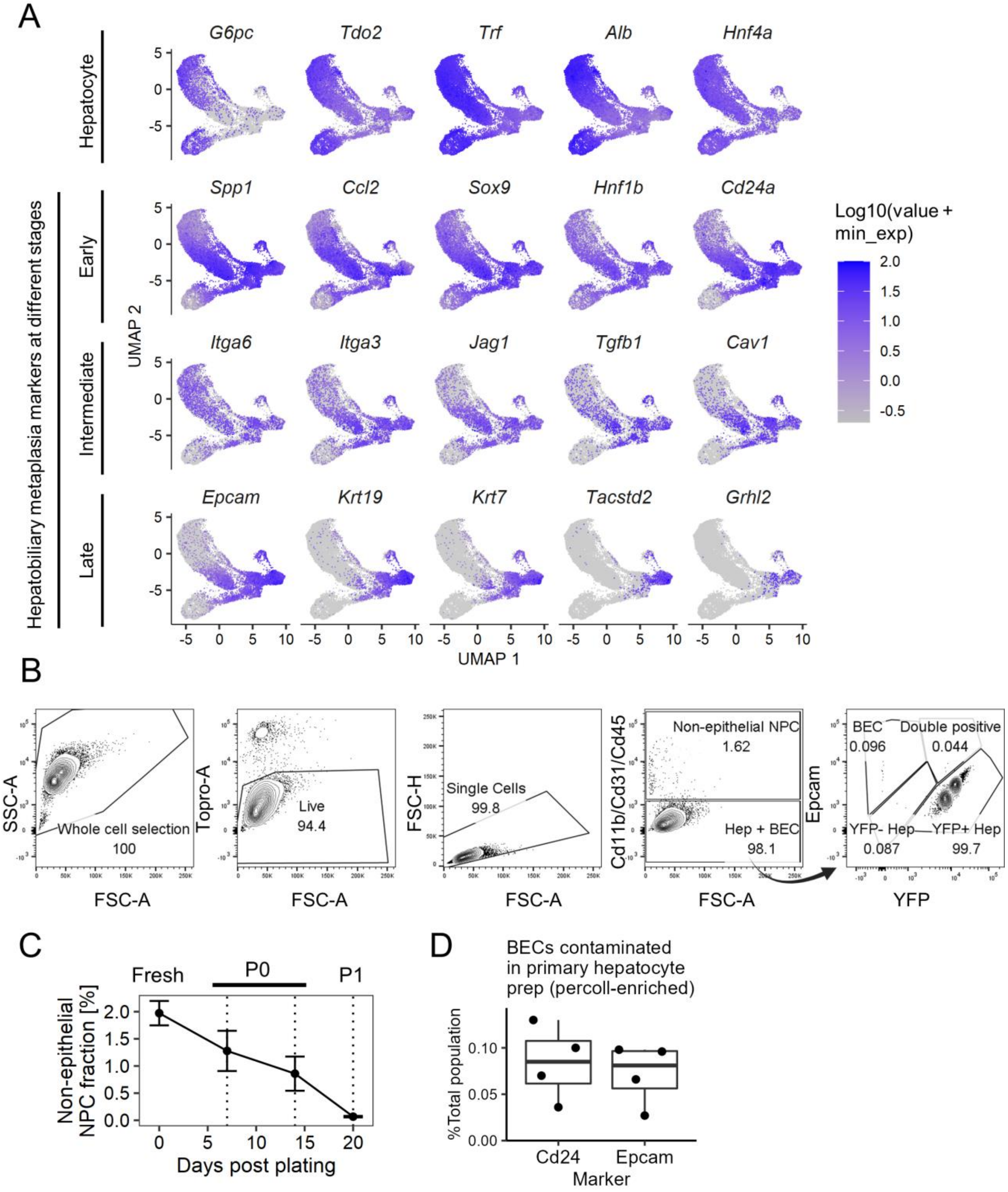


**Figure S2. Characterization of NPCs contaminated in the hepatocyte culture.**

(A) Single cell RNA-seq of whole YFP+ hepatocyte-derived cells harvested from Rosa26-LSL-YFP mice challenged with 0.1% 3,5-diethoxycarbonyl-1,4-dihydrocollidine, a toxin which can induce hepatobiliary metaplasia (n = 2 mice)[1]. Either bulk liver cells (whole Cd45-/Cd31-/Cd11b-) or YFP-sorted cells (YFP+/Cd45-/Cd31-/Cd11b-) were used. The cells with at least one YFP read count were then sorted *in silico*, and visualized by UMAP projection. Representative marker genes for the designated stages are shown.

(B) Gating strategy to determine fraction of YFP+ cells and NPCs. Epcam and Cd24 were used as BEC markers (Epcam plots are shown here as a representative example). Size selection based on FSC-A was not performed because cell sizes were largely different among cell types in the liver (leftmost panel).

(C) Time course of the fraction of non-epithelial NPC cells, which were identified as cells positive for either Cd11b, Cd31 or Cd45. Data represent mean ± SEM (n = 4 donors).

(D) Quantification of BEC contamination by flow cytometry in hepatocyte preparation by the percoll-enrichment method. Two BEC markers, Cd24 and Epcam, are used.

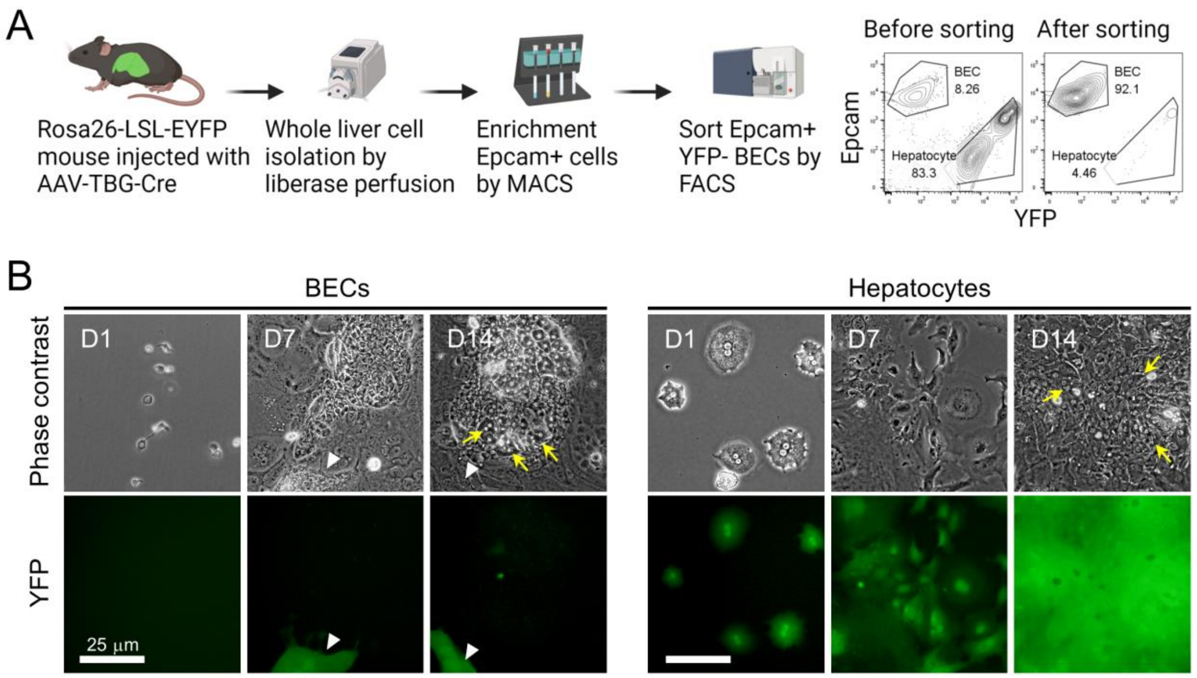


**Figure S3. Both hepatocyte-derived and BEC-derived cells spontaneously differentiate to morphologically hepatocyte-like cells.**

(A) Schematic of sorting strategy to isolate Epcam+YFP- primary BECs.

(B) Phase contrast and corresponding YFP images of BEC-derived and hepatocyte-derived proliferative cells. White arrowheads indicate hepatocyte-derived cells contaminated in the BEC culture. Yellow arrows indicate binucleated hepatocyte-like cells spontaneously differentiated from the proliferative cells in the regions with high cell density.

**Table S1. Antibodies used for immunofluorescence.**

| Antibody | Host animal | Catalog # | Dilution | Manufacturer |
|---|---|---|---|---|
| GFP | Chicken | ab13970 | 1:500 | Abcam |
| Hnf4a | Goat | sc-6556 | 1:200 | Santa Cruz |
| Hnf1b | | | 1:200 | Santa Cruz |
| Itga6 | Rat | N/A | 1:100 | V. Factor Lab |
| Epcam | Rabbit | ab71916 | 1:200 | Abcam |
| Spp1 | Goat | AF808 | 1:200 | R&D |
| Sox9 | Rabbit | ab5535 | 1:200 | Millipore |
| Krt19 | Rabbit | N/A | 1:1000 | In-house |

**Table S2. Antibodies used for flow cytometry.**

| Target | Fluorophore | Host animal | Catalog # | Dilution | Manufacturer |
|---|---|---|---|---|---|
| Epcam | BV421 | Rat | 118225 | 1:100 | BioLegend |
| Cd24 | BV421 | Rat | 101826 | 1:100 | BioLegend |
| Cd11b | PE/Cy7 | Rat | 101216 | 1:100 | BioLegend |
| Cd31 | PE/Cy7 | Rat | 102418 | 1:100 | BioLegend |
| Cd45 | PE/Cy7 | Rat | 103114 | 1:100 | BioLegend |

**Table S3. Primers used for qRT-PCR.**

| Target | Forward | Reverse |
|---|---|---|
| *Gapdh* | TCACCACCATGGAGAAGGC | GCTAAGCAGTTGGTGGTGCA |
| *Alb* | GCTGAGACCTTCACCTTCCA | CTTGTGCTTCACCAGCTCAG |
| *Asgr1* | TTGGATTGGCCTAACTGACC | GCCCATGTCCGTACCAGTTA |
| *Bmp1* | CCAGGGGCATCTTCTTGGAC | GTGCTGTCTTGGAGGGTCTC |
| *Cav1* | GCGACCCCAAGCATCTCAA | ATGCCGTCGAAACTGTGTGT |
| *Ccl2* | TAAAAACCTGGATCGGAACCAAA | GCATTAGCTTCAGATTTACGGGT |
| *Cd24a* | CTTCTGGCACTGCTCCTACC | TACTTGGATTTGGGGAAGCA |
| *Cd44* | TCGATTTGAATGTAACCTGCCG | CAGTCCGGGAGATACTGTAGC |
| *Cebpa* | CTCCCAGAGGACCAATGAAA | AAGTCTTAGCCGGAGGAAGC |
| *Cebpb* | GTTTCGGGACTTGATGCAAT | CCCGCAGGAACATCTTTAAGT |
| *Cyr61* | TAAGGTCTGCGCTAAACAACTC | CAGATCCCTTTCAGAGCGGT |
| *Epcam* | TCTACAAGGAAGAAATCAGCAAAA | CCCTCCTCAGTTCAGCACTC |
| *Fah* | CGGCGATGAAGTCATCATAA | GAGCTTCAGGCTGGTGAAAG |
| *Foxa3* | CTGGCCGAGTGGAGCTACTA | GGAGAGCTGAGTGGGTTCAA |
| *G6pc* | CTGTGCAGCTGAACGTCTGT | GAAAGTTTCAGCCACAGCAA |
| *Grhl2* | TGCCAGTGGAGAAAATCACA | TGCTCTCCATGTTGAGGATG |
| *Hes1* | GAAGCACCTCCGGAACCT | GTCACCTCGTTCATGCACTC |
| *Hnf1a* | GTCGAACATCCAGCACCTG | CCGTTGGAGTCGGAACTCT |
| *Hnf1b* | TCTCACCAGCATGTCTTCCA | AAAATGGGGTCCTTGTTGCT |
| *Hnf4a* | GCCTCAAAGCCATCATCTTC | CCGGTCGTTGATGTAATCCT |
| *Itga3* | CCTCTTCGGCTACTCGGTC | CCAGTCCGGTTGGTATAGTCATC |
| *Itga6* | TCATCCTCCTGGCTGTTCTT | GTATCGGGGAATGCTGTCAT |
| *Jag1* | GAGGCGTCCTCTGAAAAACA | TAGAAGGCTGTCACCAAGCA |
| *Krt7* | CATTGAGATCGCCACCTACC | GATAAGCTTGCCACCATTGC |

| | | |
|---|---|---|
| *Krt19* | TTGAGAGCCTGAAGGAGGAG | AATCCACCTCCACACTGACC |
| *Serpina7* | ATGCCTTTGCTGAAAGTGCT | TTCACCAATGTGTAGCACAGC |
| *Smo* | GTGCTGTCTACATGCCCAAGT | GCAACGCAGAAAGTCAGGC |
| *Sox9* | GACTCCCCACATTCCTCCTC | CCCTCTCGCTTCAGATCAAC |
| *Spp1* | GCTTGGCTTATGGACTGAGG | CGCTCTTCATGTGAGAGGTG |
| *Tacstd2* | CACGGCTCAGAGCAACTGTA | AATACCTGTGAGCCCATTGC |
| *Tat* | GTTGTCTGCCATTCCTGGAC | GCTCTGTGAATTCCACGTCA |
| *Tgfb1* | CCACCTGCAAGACCATCGAC | CTGGCGAGCCTTAGTTTGGAC |
| *Tdo2* | GGGGATCCTCAGGCTATCAT | TACCCAGTGTCTGGGAACCA |
| *Tnfrsf12a* | GTGTTGGGATTCGGCTTGGT | GTCCATGCACTTGTCGAGGTC |
| *Trf* | TAGGAGCGGAGTACATGCAA | GAGCATCTGTCTCCACCACA |
| *Ttr* | TGGACACCAAATCGTACTGG | CAGAGTCGTTGGCTGTGAAA |
| *Vim* | CGTCCACACGCACCTACAG | GGGGGATGAGGAATAGAGGCT |

### Supplemental Experimental Procedures

#### Immunofluorescence

Cells were seeded into 8-well Nunc Lab-Tek II chamber slides (Thermo Scientific) coated with collagen coating solution (Sigma), and fixed in 4% paraformaldehyde at room temperature (RT) for 15 min. Fixed cells were blocked and permeabilized in PBS with 0.3% Triton-X and 5% donkey serum for 1 h. After blocking, cells were incubated in primary antibody diluted in 5% donkey serum at RT for 1 h or overnight at 4°C. After PBS-T washes, cells were incubated with fluorescently conjugated secondary antibodies (1/300 dilution in PBS-T) and mounted with Aqua Poly/Mount (Polysciences, Inc). Slides were visualized using an Olympus IX71 inverted multicolor fluorescent microscope equipped with a DP71 camera. Primary antibodies used are listed in **Table S1**.

#### Flow cytometry and FACS

Cells were stained with antibodies listed in **Table S2** at 1/100 dilution in flow buffer (HBSS, pH 7.4, supplemented with 25 mM HEPES (Thermo), 5 mM $MgCl_2$ (MedSupply Partners), 1× Pen/Strep (Thermo), 1× Fungizone (Thermo), 1× NEAA (Thermo), 1× Glutamax (Thermo), 0.3% glucose (Sigma), 1× sodium pyruvate (Thermo)) supplemented with 40 µg/ml DNase I (hereafter called flow buffer(+)). The cells were washed twice in flow buffer(+) and centrifuged at 800 ×*g* for 1 minutes before and after each wash, then were resuspended in flow buffer(+) containing 1/1,000× TO-PRO-3 (Thermo) and analyzed using an LSR II analyzer (BD) for flow cytometry or sorted on an Aria II sorter (BD).

#### Hepatic induction

To generate spheroids, we used Aggrewell400 24-well plates (Stemcell Technologies), which were pre-treated with Anti-Adherence Rinsing Solution according to the manufacturer’s instructions (Stemcell Technologies). After trypsinization, cells were resuspeded in SHM+YAC without FBS, and the cell concentration was adjusted to 1-1.2 × $10^6$ cells/ml. Then, 1 ml of the cell suspension was added to an Aggrewell400 plate well which was pre-filled with 1 ml of SHM+YAC, and mixed thoroughly by pipetting. Then, plates were centrifuged at 100 ×*g* for 3 minutes, and placed in a $CO_2$ incubator. Following incubation for 3 days without medium change, spheroids were harvested in a 15 ml tube. After sitting for approximately 5 minutes, the supernatant was removed, and spheroids were resuspended in 5 ml SHM+YAC supplemented with 10 ng/ml mouse oncostatin M (R & D), and seeded to a 60 mm ultra-low attachable suspension culture plate (Corning). Then, spheroids were cultured on a rocking shaker at 15 rpm speed for 6-8 days. Half volume (2.5 ml) of the medium was removed and 2.5 ml fresh medium was replenished every 2 days.

#### qRT-PCR

Total RNA was extracted using a Nucleospin column (Takara) following the manufacturer’s instructions. Approximately 500 ng of RNA was reverse transcribed in 20 µl volume using a High Capacity cDNA Reverse Transcription Kit (Takara). After diluting the cDNA at a 1/20 ratio in water, qPCR was performed with primers listed in **Table S3** using SsoAdvanced SYBR reagent (Bio-Rad) and Bio-Rad CFX 384 qPCR machine.

#### RNA sequencing

In the *in vitro* experiments, RNA was isolated as described above from mouse hepCLiPs cultured under 2D and 3D cultures (n = 2 clones from 2 donor mice; one from a Rosa-LSL-EYFP mouse, and the other from a Rosa-LSL-confetti mouse). In the *in vivo* experiments, RNA was isolated from Rosa26-LSL-Cas9-EGFP mice (Katsuda et al. in prep). RNA was sent to Novogene for library preparation and high-throughput sequencing using Illumina sequencers (Hiseq 2500) to generated paired-end 150 bp data. Reads were aligned to the mouse genome (GRCm39) using STAR aligner[2]. Gene-count matrices were produced by featureCounts[3]. Gene expression levels were normalized based on the median expression level using DESeq2[4]. GSEA was performed using the fgsea (fast GSEA) R package[5]. The *in vitro* and *in vivo* RNA-seq data has been deposited to the Gene Expression Omnibus with the accession numbers GSE159885 and GSE218945, respectively.

**Single cell RNA-seq (scRNA-seq) analysis**

Data were downloaded from GEO (accession number GSE157698). Using R Seurat package, Seurat objects were created with arguments “min.cells = 3, min.genes = 200”. The cells were further filtered with a “subset” function with arguments “subset = nFeature_RNA > 200 & nFeature_RNA < 4000 & percent.mt < 0.25 & percent.yfp > 0”, which filtered out YFP-negative cells. UMAP was projected using a monocle3 package with “plot_cells” function and an argument “scale_to_range = TRUE”.

**Intrasplenic transplantation of hepCLiPs**

Two-dimensional cultured cells were harvested using TrypLE (Thermo). Briefly, hepCLiPs grown in a 10 cm plate were treated with 2 ml TrypLE at 37 °C for approximately 10 min, and harvested with 5 ml 10%FBS-DMEM. Three-dimensional cultured spheres were collected in 15 ml tubes by gravity for approximately 1 min. After washed twice in PBS (collected by gravity), the spheres were dissociated with TrypLE Express or Accutase (Sigma) supplemented with 40 µg/ml DNase I at 37 °C for 25-45 minutes with intermittent inversion. Following further dissociation with gentle pipetting with P1000 tips, the cells were resuspended in 5 ml 10%FBS-DMEM supplemented with 0.04 mg/ml DNase. Harvested cells were then filtered with a 35 µm cell strainer-equipped FACS tubes (BD) and collected by centrifugation at 1,500 rpm (approximately 480 ×*g*) for 3 min. For hepCLiP_2D and hepCLiP_3D cells harvested with TrypLE, the cells were resuspended in HBSS at $1 \times 10^7$ cells/ml. Then, a small incision was made on the left flank of the mice, the spleen was extracted and held with tweezers, and $5 \times 10^5$ cells/50 µl were injected into the spleen of FRG mice. For hepCLiP_3D cells harvested with Accutase, the cells were resuspended in HBSS at $5 \times 10^6$ cells/ml, and $2.5 \times 10^5$ cells/50 µl were injected into the spleen of FRG mice. All the transplantation procedures were conducted by the Penn Xenograft Core. The FRG mice were maintained under nitisinone cycles following the guideline by Yecuris™, the animal supplier, and all animals were maintained by the Penn Xenograft Core.

**Imaging of the repopulated livers**

Three months after transplantation, the livers were harvested following blood removal by portal vein perfusion of ~30 ml HBSS. The extracted livers were imaged using a Leica stereo microscope at 7.1× magnification. Repopulation rates were roughly estimated using the gross fluorescent images analyzed on ImageJ version 1.53u. The CFP+ nodules were manually counted.
